# Par protein localization during the early development of *Mnemiopsis leidyi* suggests different modes of epithelial organization in Metazoa

**DOI:** 10.1101/431114

**Authors:** Miguel Salinas-Saavedra, Mark Q Martindale

**Affiliations:** The Whitney Laboratory for Marine Bioscience, and the Department of Biology, University of Florida, 9505 N, Ocean Shore Blvd, St. Augustine, FL 32080-8610, USA

**Author notes:** Correspondence; (MS-S) (MQM). Centre for Chromosome Biology, Bioscience Building, National University of Ireland Galway, Galway, Ireland (Miguel Salinas-Saavedra).

## Abstract

In bilaterians and cnidarians, embryonic and epithelial cell-polarity are regulated by the interactions between Par proteins, Wnt/PCP signaling pathway, and cell-cell adhesion. Par proteins are highly conserved across Metazoa, including ctenophores. But strikingly, ctenophore genomes lack components of the Wnt/PCP pathway and cell-cell adhesion complexes; raising the question if ctenophore cells are polarized by mechanisms involving Par proteins. Here, by using immunohistochemistry and live-cell imaging overexpression of specific mRNAs, we describe for the first time the subcellular localization of selected Par proteins in blastomeres and epithelial cells during the embryogenesis of the ctenophore *Mnemiopsis leidyi*. We show that these proteins distribute differently compared to what has been described for other animals, even though they segregate in a host-specific fashion when expressed in cnidarian embryos. This differential localization might be related to the emergence of different junctional complexes during metazoan evolution. Data obtained here challenge the ancestry of the apicobasal cell polarity and raise questions about the homology of epithelial tissue across the Metazoa.

## INTRODUCTION

The emergence of epithelial tissues was arguably one of the most critical events in animal evolution. In bilaterians and cnidarians, a ‘true-epithelium’ is classically defined as a group of polarized cells joined by belt-like cell-cell junctions and supported by a basement membrane. Epithelial cells are polarized along the apical-basal axis and form a planar sheet of cells that can undergo subsequent morphogenesis^1–5^. While the asymmetric cortical distribution of the Wnt Planar Cell Polarity (PCP) pathway components polarizes the cells along the tissue plane, the asymmetric cortical distribution of Par system components polarizes the cells along the apical-basal axis^2,3,5–19^. These mechanisms that organize cell-polarity are highly conserved in all animals that have been studied and most likely been present in the most recent common ancestor (MRCA) of Cnidaria and Bilateria^3,5,16,18,20–23^. However, little is known outside of those two clades (Figure 1A).

**Figure 1.**
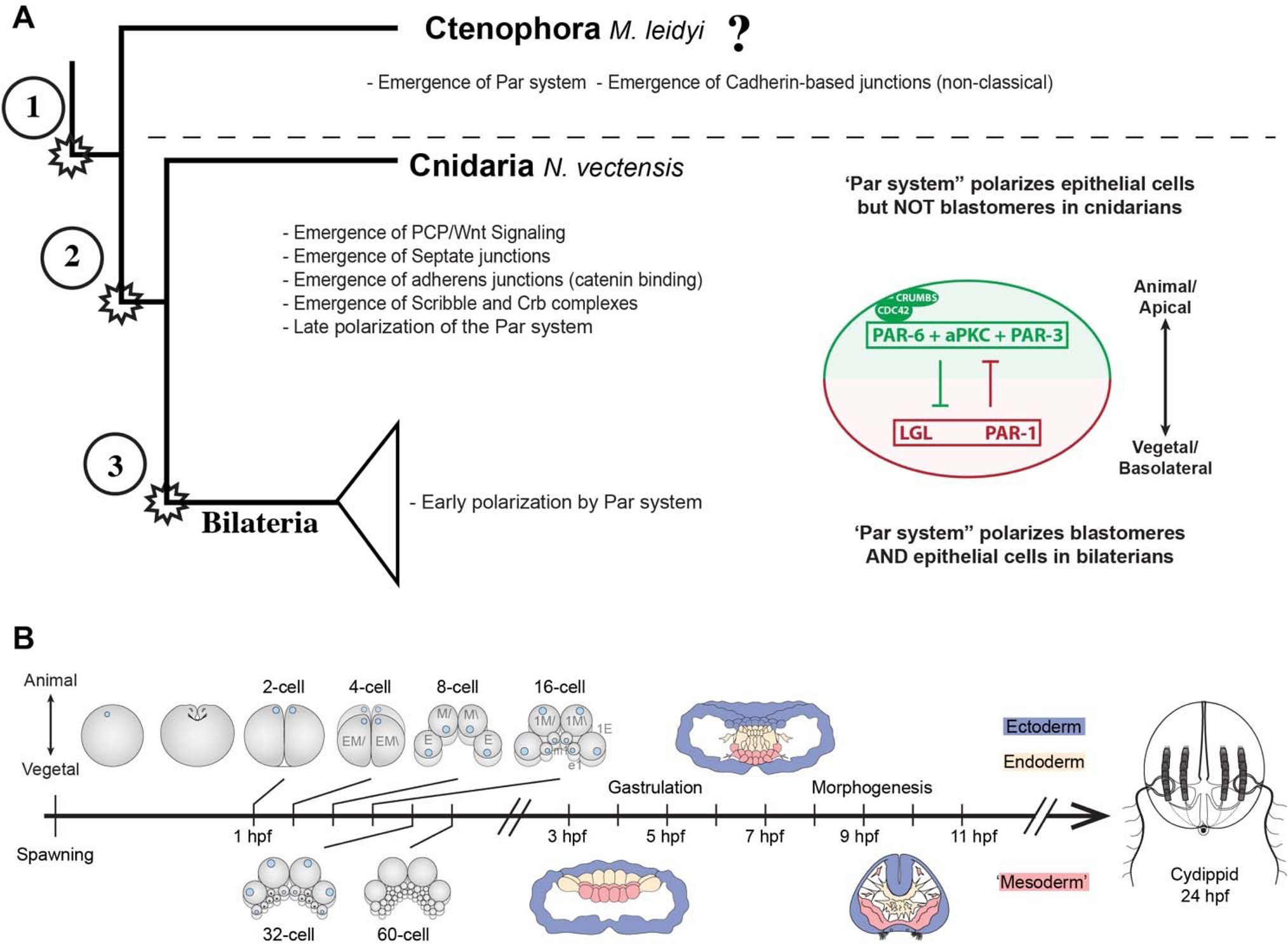
Evolution of cell polarity components during animal evolution. A) Three major evolutionary steps (left side) that might have changed the organization of cell polarity in the Metazoa. The diagram (right side) depicts the subcellular asymmetric localization of Par proteins in Cnidaria and Bilateria. However, there are no previous descriptions available for ctenophore cells. B) The stereotyped early development of *M. leidyi*.

Interestingly, ctenophores or comb jellies, whose position at the base of metazoans three is still under debate^24–28,29,30^, do not have the genes that encode the components of the Wnt/PCP pathway in their genomes^26^. Thus, the study of the subcellular organization of the Par system components in ctenophores and their potential role in apical-basal polarity is important to understand the evolution of tissue organization in Metazoa.

The asymmetric localization of the Crumbs (Crb) complex, (e.g. Crb/Pals1/Patj), the Par/aPKC complex (e.g. Par-3/aPKC/Par-6), and the Scribble complex (e.g. Scribble/Lgl/Dlg) in the cortex of bilaterian and cnidarian cells maintains epithelial integrity by stabilizing cell-cell junctions^4,5,20,22,23^. When the Par/aPKC complex is activated by Cdc42, they bind to the apical cortex via the Crb complex^4,31–33^, and recruits and stabilizes E-cadherin^4,22,34–36^. E-cadherin, in turn, sequesters ß-catenin from the cytoplasm that binds to alpha-catenin (forming the Cadherin-Catenin complex; CCC) and links this protein complex to the actin cytoskeleton, thus stabilizing Adherens Junctions (AJs)^1,23,35,37–41^. The maturation of AJs is essential for the maintenance of the Par/aPKC complex localization at the apical cortex that displaces members of the Scribble complex and Par-1 to basolateral localizations associated with Septate Junctions (SJs)^23,42–50^. Therefore, the regulation of AJs is critical for the maintenance of epithelial polarity and its functional regulation of epithelial physiology^21,22,36,51–53^.

This mechanism is deployed in metazoan cells to establish embryonic and epithelial cell polarity during early development and is critical for axial organization^5,15,17,21,22,36,54–69^. In most bilaterian cells, symmetry breaking is mediated by the Par system set up maternally during the earliest cleavage stages^5,60,70,71^. The asymmetric localization of Par proteins ensures the proper partitioning of maternal determinants into distinct daughter cells during the stereotyped cleavage program in bilaterians^3,15,55,60^. However, this is not the case in cnidarians (e.g. corals, sea anemones and “jellyfish”) that do not possess a stereotyped cell division pattern^5,72,73^. Intriguingly, ctenophores (comb jellies like *Mnemiopsis leidyi*), that are thought to be an even more ancient group of animals^24,26,28^, have polarized epithelial cells and develop under a highly determined and stereotyped developmental program^74–77^ (Figure 1B). Therefore, the description of Par protein localization will give insights on how cell polarity is deployed during these early stages.

Components of the Par system are unique to, and highly conserved, across Metazoa, including placozoans, poriferans, and ctenophores^20, 23^. But strikingly, ctenophore genomes do not have many of the crucial regulators present in other metazoan genomes^23, 48^. For example, none of the components of the Crb complex, a Scribble homolog, or SJs, are present, and the cytoplasmic domain of cadherin lacks the crucial biding sites to catenins that interact with the actin cytoskeleton^23^. These data raise the question of whether or not ctenophore cells are polarized by mechanisms involving the apicobasal cell polarity mediated by Par proteins and question the homology of epithelial tissue organization across the Metazoa. Here, by using antibodies raised to specific ctenophore proteins and confirmed by live-cell imaging of injected fluorescently labeled mRNAs, we describe for the first time the subcellular localization of selected components of the Par system during the development of the ctenophore *M. leidyi*. Data obtained here challenge the ancestry of the apicobasal cell polarity module and raise questions about the classical definition of ‘true-epithelial’ tissue as a conserved trait in all metazoans.

## RESULTS

### Antibody Specificity

Genome searches of *M. leidyi* showed that there is only a single copy for both Par-6 (*Ml*Par-6) and Par-1 (*Ml*Par-1) genes and they express through development in the transcriptome of *M. leidyi*^78, 79^. We used rabbit polyclonal affinity-purified antibodies (Bethyl labs, Inc) designed against *M. leidyi Ml*Par-6 and *Ml*Par-1 to determine their spatial and temporal expression at different developmental stages (see Materials and Methods for target sequence information). Western blots of *M. leidyi* adult extracts (Figure 2A) showed the specificity of each antibody that recognized a single band for *Ml*Par-6 (predicted size 33.3 KD; Figure 2A) and *Ml*Par-1 (predicted size 84.7 KD; Figure 2A). Pre-adsorption of the *Ml*Par-6 and *Ml*Par-1 antibody with a tenfold molar excess of the respective antigenic peptide (used to generate and affinity purify the antibodies) resulted in the elimination of the appropriate-sized single band for *Ml*Par-6 and *Ml*Par-1 (Figure 2A). In addition, pre-adsorption experiments were performed by whole-mount immunohistochemistry to test the specificity of the *Ml*Par-6 and *Ml*Par-1 antibodies. The staining pattern was strongly mitigated in early embryos when we used pre-incubated antibodies (Figure 2B). Thus, both antibodies are specific to their intended targets and provide robust reagents to determine the subcellular localization of these proteins during *M. leidyi* embryogenesis.

**Figure 2.**
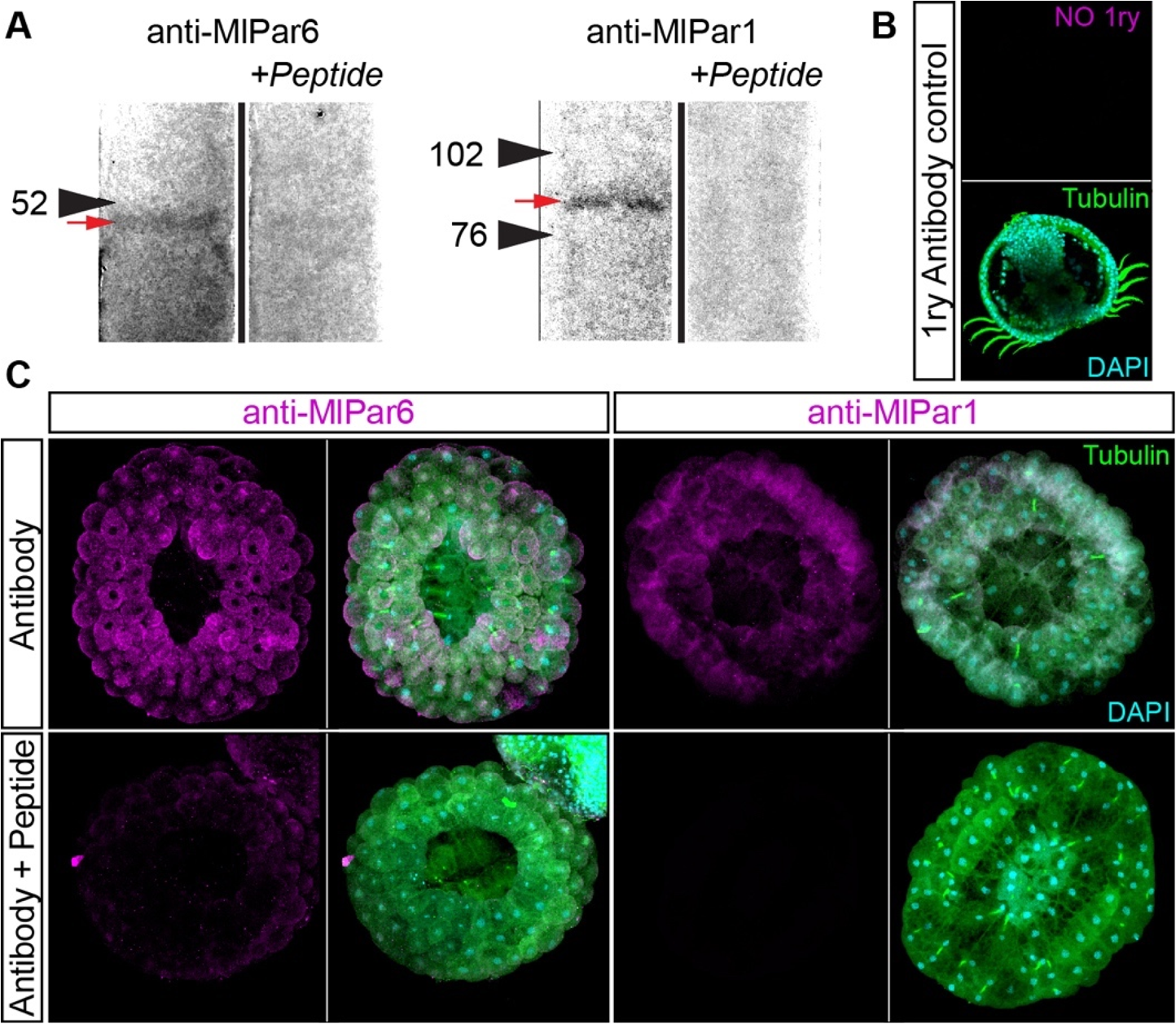
Specificity of *M. leidyi* antibodies as tested by pre-adsorption experiments. A) Western blots of *M. leidyi* adult tissue extracts using specific antibodies against *Ml*Par-6 and *Ml*Par-1. Pre-adsorption of the antibodies with a tenfold excess of the antigen peptide resulted in the elimination of the staining of the single *Ml*Par-1 band. Arrowheads indicate the molecular weight in KD. A red arrow indicates the band recognized by the antibody for each protein. See Supplementary Figure 1 for the full lanes and input lysate staining. B) Negative control without the rabbit primary antibody shows that there is no inherent autofluorescence. C) Whole-mount immunohistochemistry pre-adsorption experiments show that the staining pattern was strongly mitigated in early embryos when pre-incubated antibodies against *Ml*Par-6 and *Ml*Par-1 with the respective peptide.

### *Ml*Par-6 gets localized to the apical cortex of cells during early *M. leidyi* development

We characterized the subcellular localization of the *Ml*Par-6 protein during early *M. leidyi* development by using our specific *Ml*Par-6 antibody. In all of over 100 specimens examined, *Ml*Par-6 expression polarizes to the animal cortex (determined by the position of the zygotic nucleus; Figure 3A) of the single cell zygote and to the apical (animal) cell cortex during every cleavage stage (Figure 3). At the cortex, *Ml*Par-6 localizes perpendicular to the cleavage furrow in cell-contact-free regions facing the external media (Figure 3). As cleavage ensues, *Ml*Par-6 becomes localized to the position of cell-cell contacts between blastomeres until 3 hpf (Figure 3B-C). During gastrulation (3-7 hpf; Figures 4 and 5), *Ml*Par-6 is not localized in cells undergoing cellular movements including the oral (4 hpf; Figure 4) and aboral ectoderm (5-6 hpf; Figure 4) undergoing epibolic movements, syncytial endoderm, and mesenchymal ‘mesoderm.’ However, this protein remains apically polarized in ‘static’ ectodermal cells remaining at the animal pole (blastopore) and vegetal pole (4-7 hpf; Figures 4 and 5). By the end of gastrulation (8-9 hpf; Figure 5), *Ml*Par-6 becomes localized asymmetrically to the apical cortex of the ectodermal epidermal cells and the future ectodermal pharyngeal cells that start folding inside the blastopore (Figure 5). Interestingly, we do not observe a clear cortical localization in later cydippid stages, and the antibody signal is weaker after 10 hpf in juveniles (Figure 6). Contrary to expectations, at these later stages, *Ml*Par-6 is cytosolic and does not localize in the cortex of epidermal cells, and a few epithelial and mesenchymal cells showed nuclear localization (Figure 6A). Thereafter, *Ml*Par-6 remains cytosolic in all scored stages up to 24 hpf. Cytosolic and nuclear localization of Par-6 has been reported in other organisms when the polarizing roles of this protein are inactive^808182^. Thus, our data suggest that *Ml*Par-6 does not play a role in cell adhesion or cell polarity during juvenile cydippid stages.

**Figure 3.**
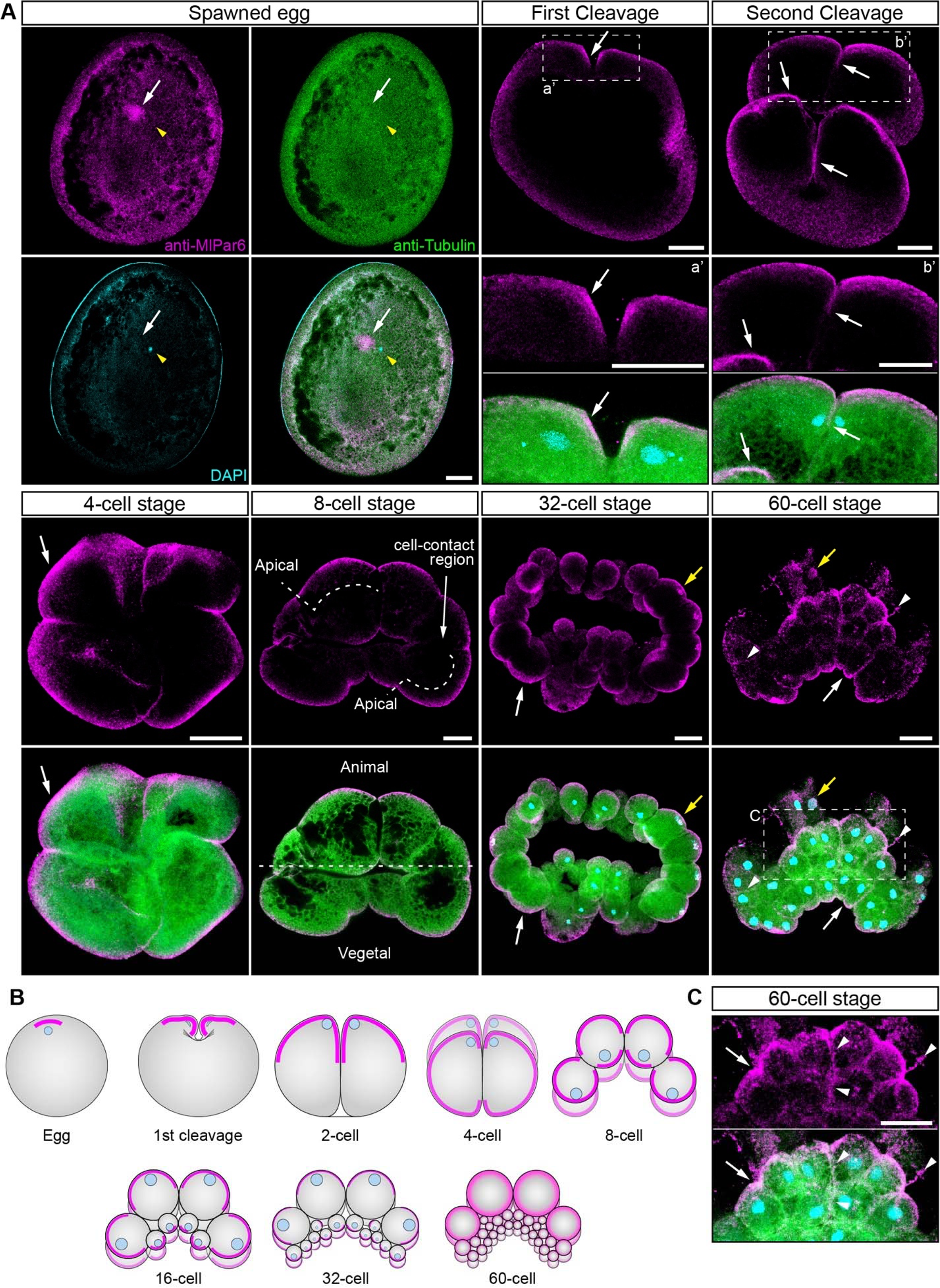
*Ml*Par-6 protein localizes asymmetrically in the cell cortex of the eggs and in the cell-contact-free regions of cleavage stages. A) Immunostaining against *Ml*Par-6 during cleavage stages of *M. leidyi* development. *Ml*Par-6 protein localizes to the apical cortex (white arrows) until the 60-cell stage where its signal was detected in regions of cell-contact (white arrowhead). Images are 3D reconstructions from a z-stack confocal series. The 8-cell stage corresponds to a single optical section. *Ml*Par-6 protein localizes to the apical cortex of the cells but is absent from cell-contact regions. Orientation axes are depicted in the Figure. a’ and b’ are magnifications of the sections depicted in the first and second cleavage, respectively. B) Diagram depicting the cortical localization of *Ml*Par-6. Animal pole is to the top as depicted in A. C) Magnification of the region c depicted in 60 cell-stage. Morphology is shown by DAPI and Tubulin immunostainings. Yellow arrowhead indicates the zygotic nucleus. Yellow arrows indicate nuclear localization. Homogeneous cytosolic staining was digitally reduced to highlight cortical localizations. See Supplementary Figure 2 for a general picture of the cleavage stages. Scale bars: 20 µm.

**Figure 4.**
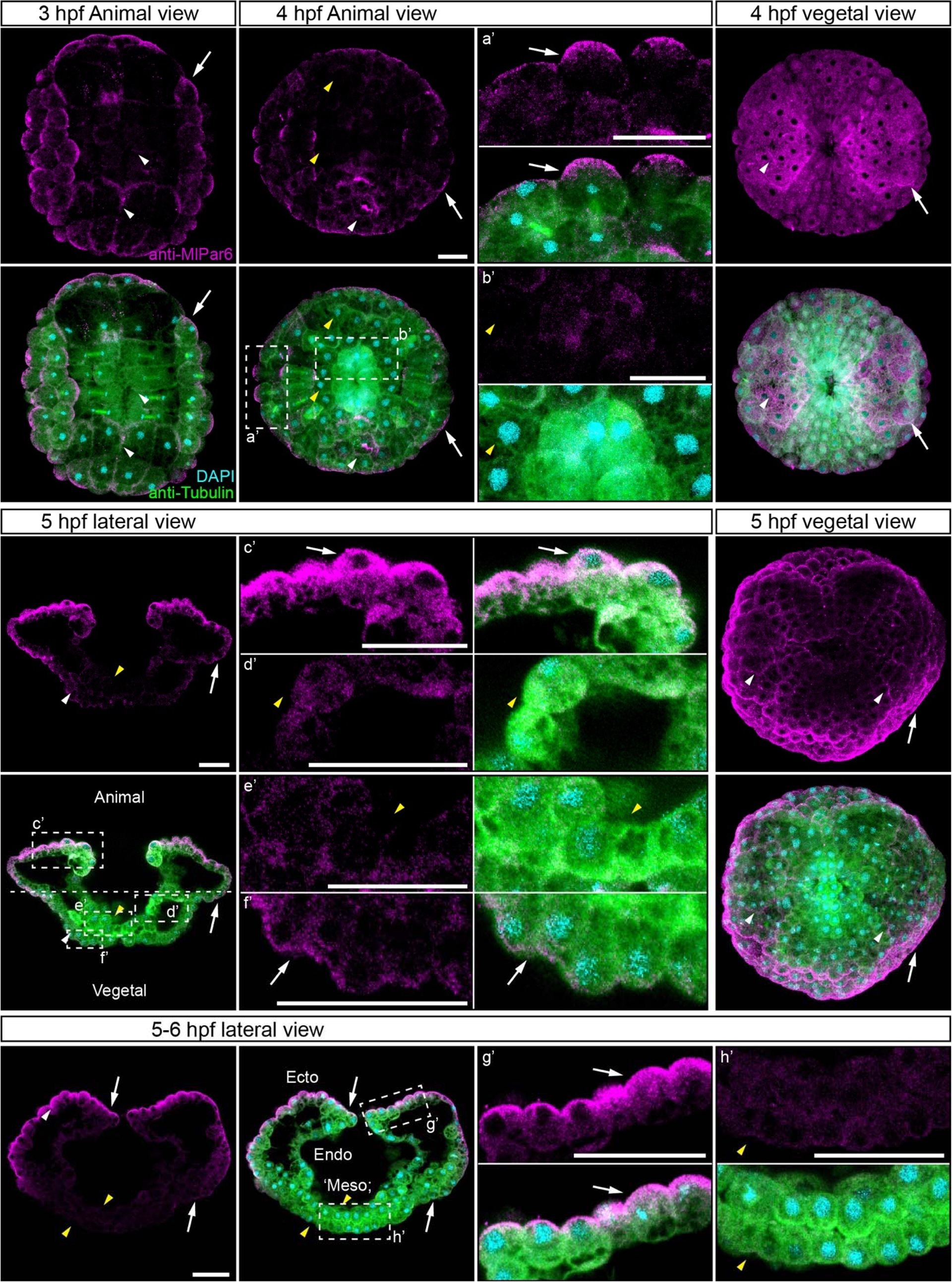
*Ml*Par-6 protein remains apically polarized in ‘static’ ectodermal cells during *M. leidyi* gastrulation. Immunostaining against *Ml*Par-6 during early gastrulation stages of *M. leidyi* development. *Ml*Par-6 protein localizes to the apical cortex (white arrows) but it is not localized in cells undergoing cellular movements. Images are 3D reconstructions from a z-stack confocal series. 5-6 hpf, *Ml*Par-6 protein localizes to the apical cortex of the ectodermal cells (Ecto) but is absent from endodermal (Endo) and ‘mesodermal’ (‘Meso’) cells. a’ to h’ correspond to magnifications of the regions depicted for each stage. Axial orientation is depicted with the animal pole to the top. Morphology is shown by DAPI and Tubulin immunostainings. White arrowheads indicate *Ml*Par-6 protein in regions of cell-contact. Yellow arrowheads indicate the absence of cortical localization. Homogeneous cytosolic staining was digitally reduced to highlight cortical localizations. Scale bars: 20 µm.

**Figure 5.**
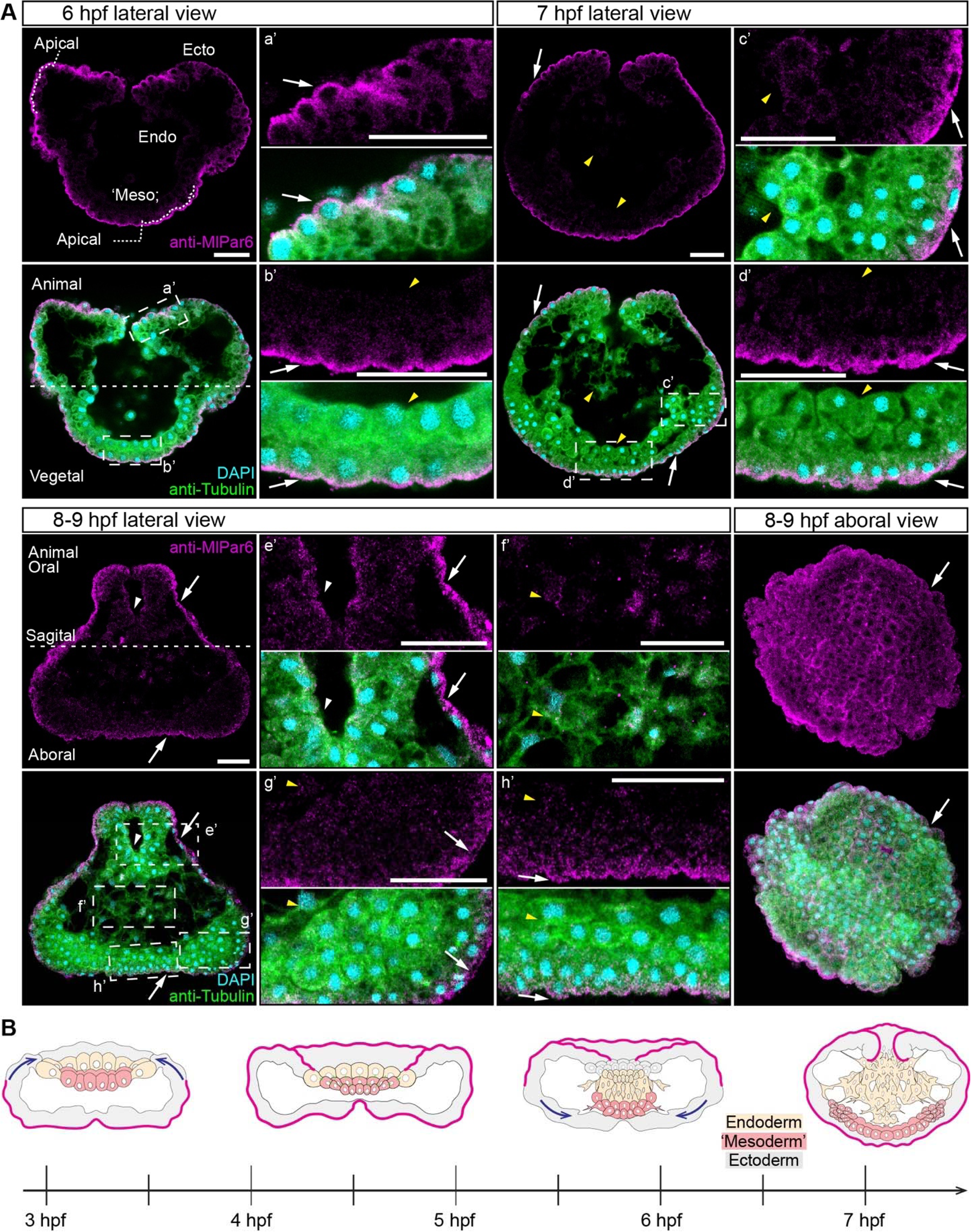
*Ml*Par-6 protein localization during late *M. leidyi* gastrulation. A) Immunostaining against *Ml*Par-6 during late gastrulation stages of *M. leidyi* development. *Ml*Par-6 protein localizes to the apical cortex of the ectodermal cells (Ecto) but is absent from endodermal (Endo) and ‘mesodermal’ (‘Meso’) cells. *Ml*Par-6 protein localizes to the apical cortex of the ectoderm (white arrows) until 9 hpf. Images are 3D reconstructions from a z-stack confocal series. 6 hpf embryo corresponds to a single optical section from a z-stack confocal series. a’ to h’ correspond to the magnifications of the regions depicted for each stage. The axial orientation in each panel is animal pole up. B) Diagram depicting the cortical localization of *Ml*Par-6 (magenta). Ectoderm is colored in grey. Endoderm and ‘mesoderm’ are colored in yellow and red, respectively. Blue arrows depict gastrulation movements. For simplicity, most of the cell boundaries are not depicted. The animal pole is to the top as depicted in A. Morphology is shown by DAPI and Tubulin immunostainings. White arrowhead indicates *Ml*Par-6 protein in pharyngeal cells. Yellow arrowheads indicate the absence of cortical localization. Homogeneous cytosolic staining was digitally reduced to highlight cortical localizations. Scale bars: 20 µm.

**Figure 6.**
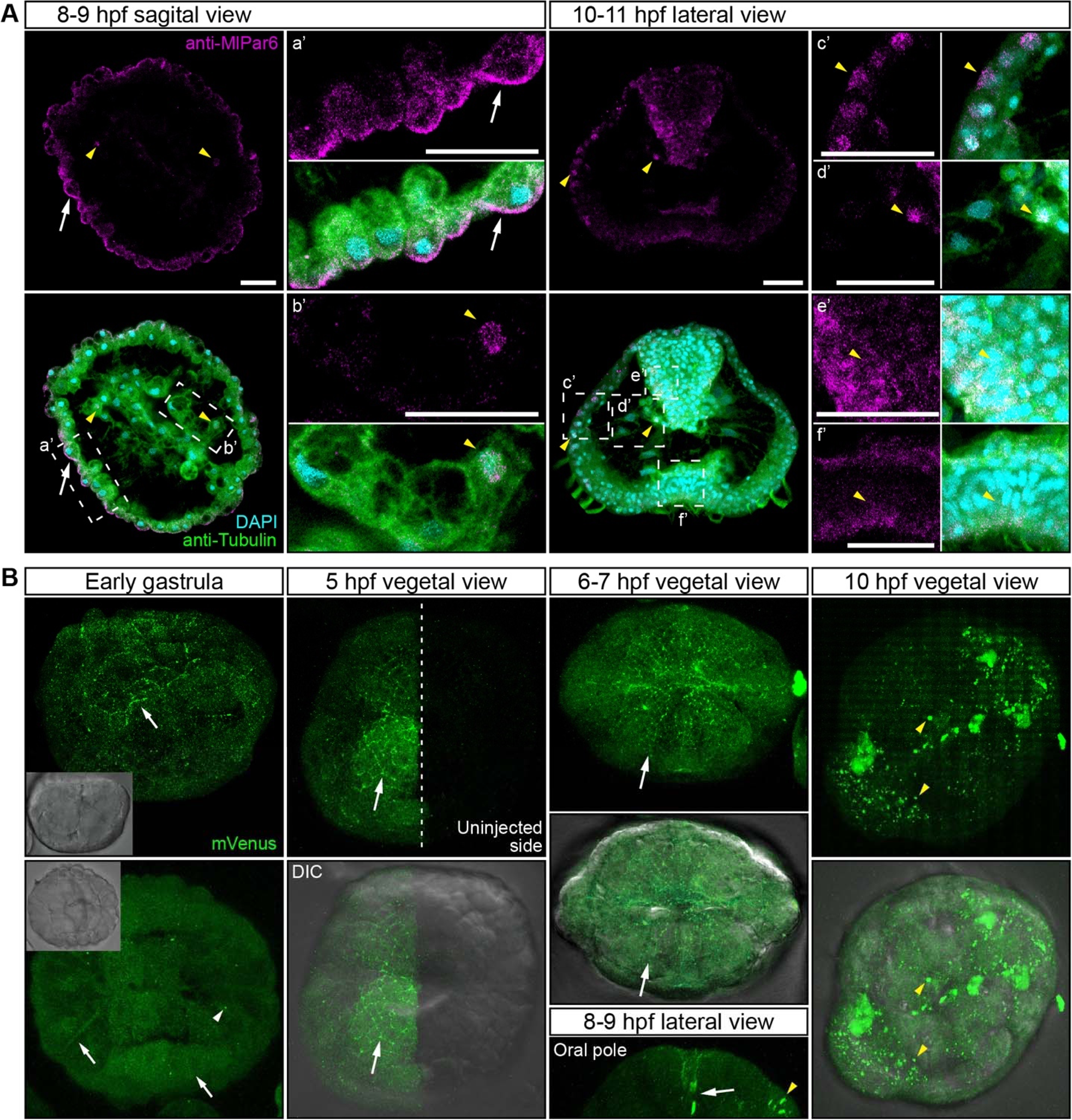
*Ml*Par-6 protein does not localize to the cortex of epidermal cells after 10 hpf during *M. leidyi* development. A) Immunostaining against *Ml*Par-6 shows that *Ml*Par-6 protein localizes to the apical cortex (white arrows) until 9 hpf but it is not cortically localized after 10 hpf. 8-9 hpf images are single optical sections from a z-stack confocal series. 10-11 hpf images are 3D reconstructions from a z-stack confocal series. a’ to f’ correspond to the magnifications of the regions depicted for each stage. Orientation is depicted in the Figure: Animal/oral pole is to the top. Morphology is shown by DAPI and tubulin immunostainings. Yellow arrowheads indicate nuclear localization. See Supplementary Figure 3 for a later stage. B) *In vivo* localization of *Ml*Par6-mVenus during different stages of *M. leidyi* development. The overexpression of *Ml*Par6-mVenus protein displays similar patterns observed with the antibody staining against the same protein, with no cortical localization was observed after 10 hpf. All images are 3D reconstructions from a z-stack confocal series. Orientation is animal/oral pole to the top. Morphology is shown by DIC microscopy. White arrows indicate *Ml*Par6-mVenus protein cortical localization. Yellow arrowheads indicate the absence of cortical localization and cytosolic aggregation. Homogeneous cytosolic staining was digitally reduced to highlight cortical localizations. Scale bars: 20 µm.

Similar results were obtained when we overexpressed the mRNA encoding for *Ml*Par-6 fused to mVenus (*Ml*Par-6-mVenus) and recorded the *in vivo* localization of the protein in *M. leidyi* embryos (Figure 6B). *M. leidyi* develops rapidly with cell cycle times of 15 minutes between cleavages, and therefore, it was impossible at this time to observe the *in vivo* localization of this protein in early blastomeres because in these experiments, the translated *Ml*Par-6-mVenus was observed approximately 4 hours post injection into the uncleaved egg. During gastrulation, *Ml*Par-6-mVenus localizes to the apical cell cortex and displays enrichment at the level of cell-cell contacts (Figure 6B). However, as we observed by antibody staining, this cortical localization is no longer observable during the cell movements associated with gastrulation and *Ml*Par-6-mVenus remains cytosolic (Figure 6B). After 8 hpf, *Ml*Par-6-mVenus localizes to the apical cortex of ectodermal epidermal and pharyngeal cells but is not observable in any other internal tissue (Figure 6B). After 10 hpf, *Ml*Par-6-mVenus remains in the cytosol and no cortical localization was detectable (Figure 6B), confirming the antibody observations presented above.

### *Ml*Par-1 remains cytoplasmic during early *M. leidyi* development

In bilaterians and cnidarians, the apical localization of *Ml*Par-6 induces the phosphorylation of *Ml*Par-1 displacing this protein to basolateral cortical regions^4,5,21,22^. Using our specific *Ml*Par-1 antibody, we characterized the subcellular localization of the *Ml*Par-1 protein during the early *M. leidyi* development (Figures 7, 8, and 9). Even though *Ml*Par-1 appears to be localized in the cortex at the cell-contact regions of early blastomeres (Figure 7) and gastrula stages (Figure 8), this antibody signal was not clear enough to be discriminated from the cytosolic distribution, possibly due to edge effects. Nevertheless, and strikingly, *Ml*Par-1 remains as punctate aggregations distributed uniformly in the cytosol, and in some cases, co-distributes with chromosomes during mitosis (Figures 7, 8, and 9). We did not observe asymmetric localization of *Ml*Par-1 in the cell cortex of *M. leidyi* embryos at any of the stages described above for *Ml*Par-6.

**Figure 7.**
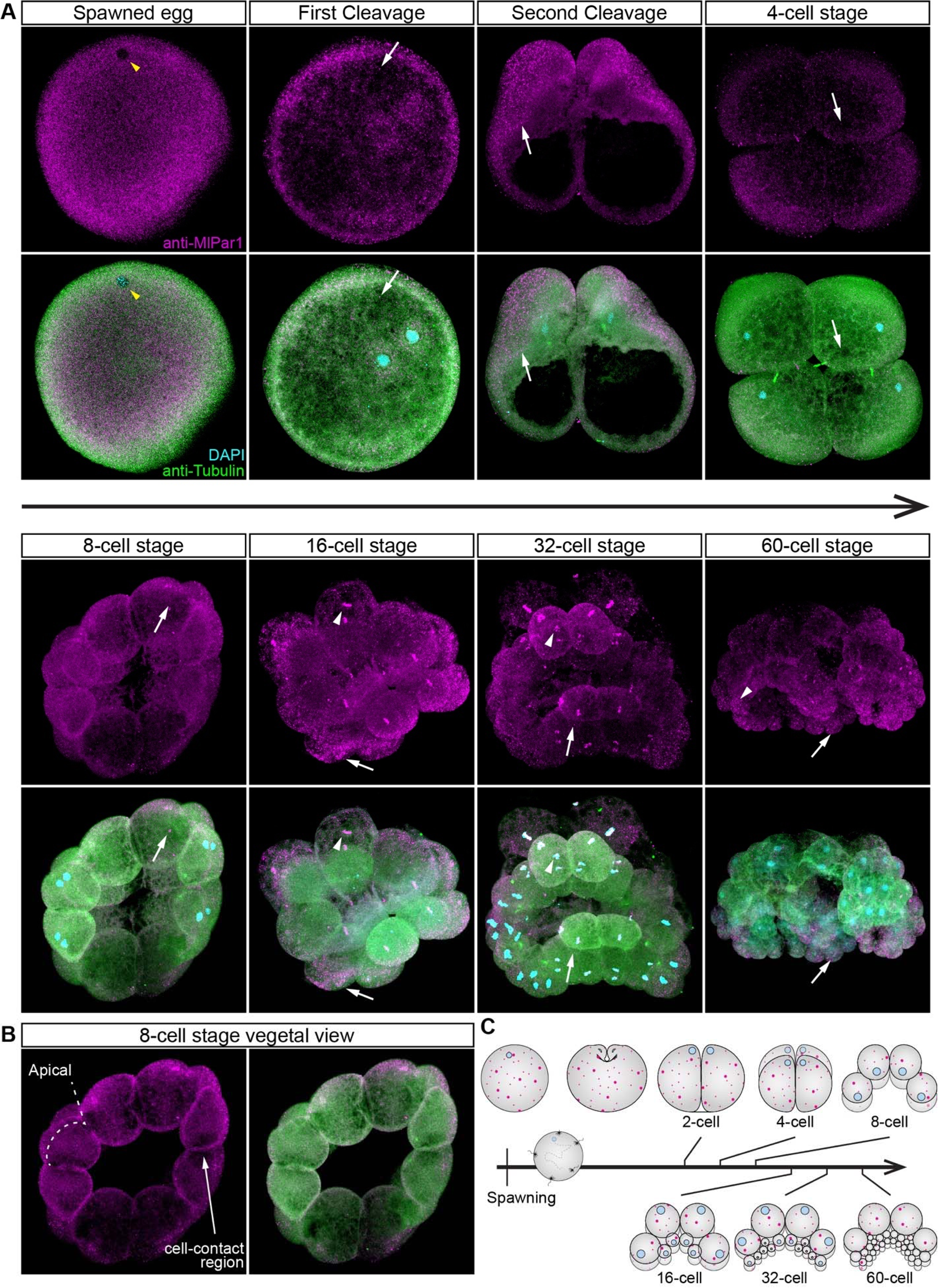
*Ml*Par-1 protein remains cytoplasmic during early cleavage stages. A) Immunostaining against *Ml*Par-1 during cleavage stages of *M. leidyi* development. *Ml*Par-1 protein appears as punctate aggregations distributed uniformly in the cytosol (white arrows). Images are 3D reconstructions from a z-stack confocal series. B) A single optical section from the 8-cell stage z-stack confocal series shown in A. *Ml*Par-1 appears to be localized in the cortex at the cell-contact regions but this antibody signal was similar to its cytosolic distribution. C) Diagram depicting the localization of *Ml*Par-1. The animal pole is oriented towards the top. Morphology is shown by DAPI and tubulin immunostainings. Yellow arrowhead indicates the zygote nucleus.

**Figure 8.**
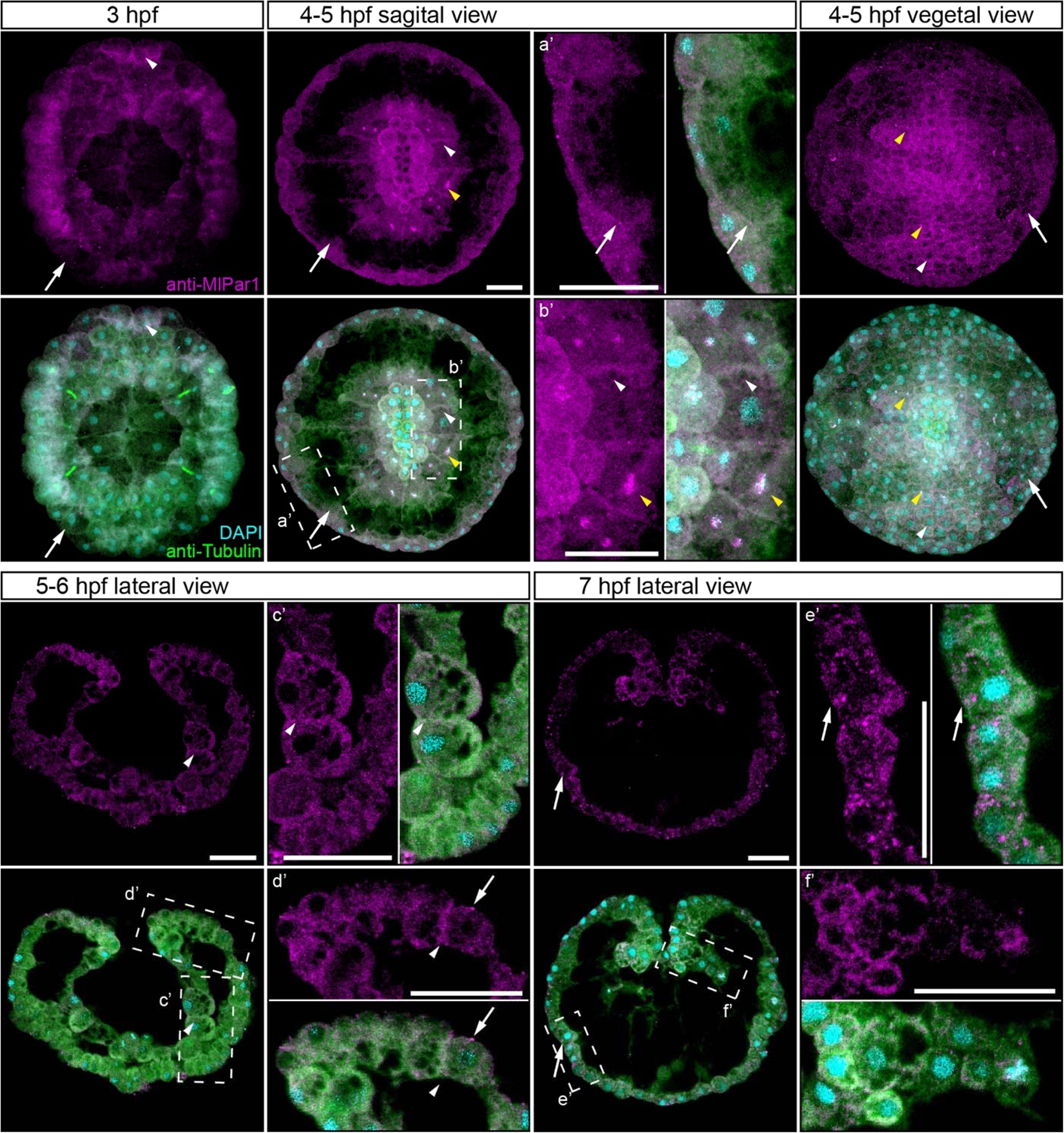
*Ml*Par-1 protein remains cytoplasmic during *M. leidyi* development. Immunostaining against *Ml*Par-1 between 3 hpf and 7 hpf. *Ml*Par-1 protein remains as punctate aggregations distributed uniformly in the cytosol (white arrows). *Ml*Par-1 appears to be localized in the cortex at the cell-contact regions (white arrowheads) but this antibody signal was similar to its cytosolic distribution. Images are 3D reconstructions from a z-stack confocal series. a’ to f’ correspond to the magnifications of the regions depicted for each stage. Orientation axes are depicted in the figure. Morphology is shown by DAPI and tubulin immunostainings. Yellow arrowheads indicate nuclear localization. The animal pole is towards the top. Scale bars: 20 µm.

**Figure 9.**
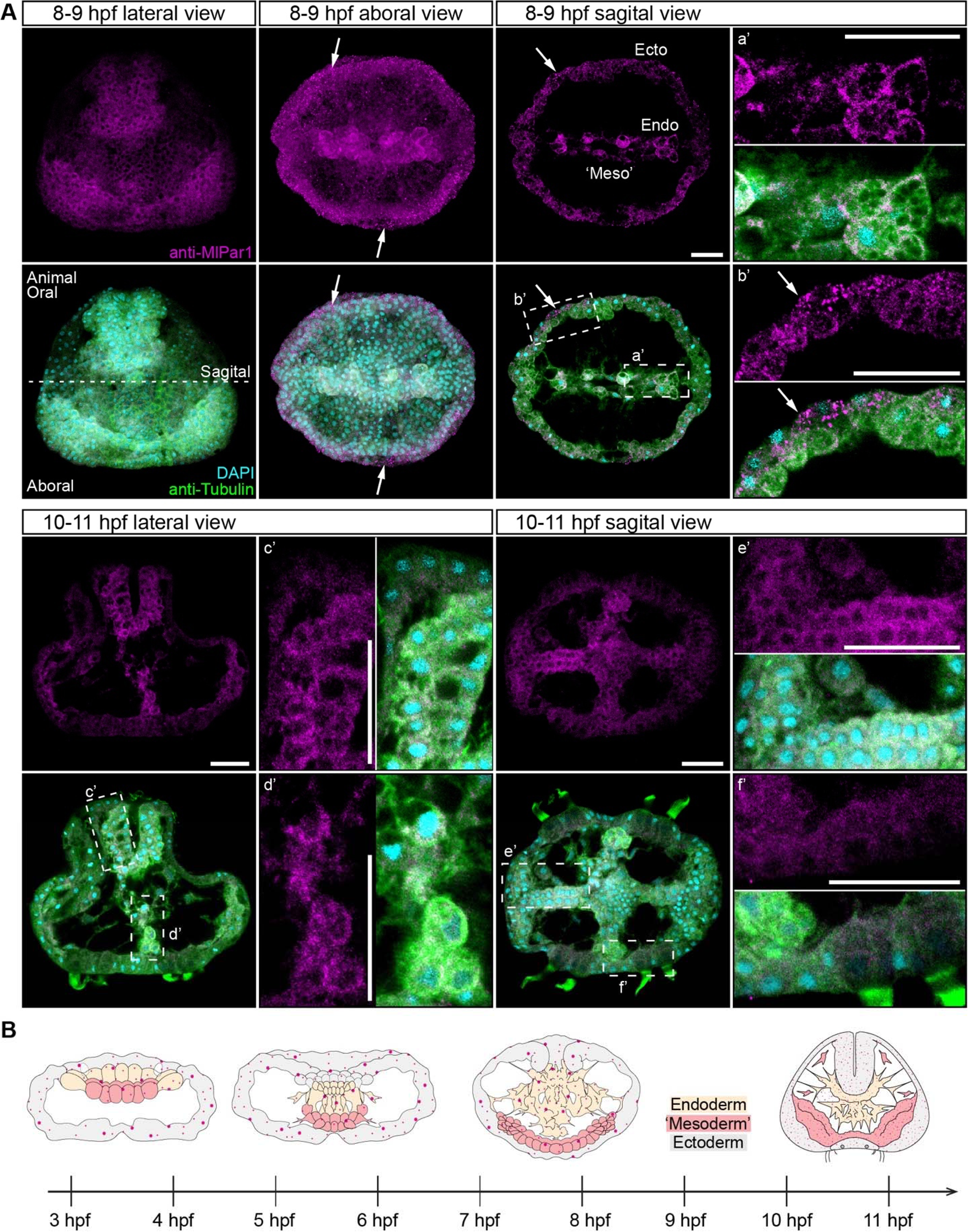
*Ml*Par-1 protein remains cytoplasmic during *M. leidyi* development between 8 hpf and 11 hpf. *Ml*Par-1 protein remains as punctate aggregations distributed uniformly in the cytosol (white arrows). Images are 3D reconstructions from a z-stack confocal series. Sagittal view of an 8-9 hpf embryo corresponds to a single optical section from a z-stack confocal series. *Ml*Par-1 protein remains cytoplasmic in ectodermal cells (Ecto), endodermal (Endo), and ‘mesodermal’ (‘Meso’) cells. a’ to f’ correspond to the magnifications of the regions depicted for each stage. B) Diagram depicting the localization of *Ml*Par-1 (magenta). Ectoderm is colored in grey. Endoderm and ‘mesoderm’ are colored in yellow and red, respectively. For simplicity, most of the cell boundaries are not depicted. Morphology is shown by DAPI and tubulin immunostainings. Yellow arrowheads indicate nuclear localization. The animal pole is to the top. Scale bars: 20 µm.

We observed these similar results *in vivo* when we overexpressed the mRNA encoding for *Ml*Par-1 fused to mCherry (*Ml*Par-1-mCherry) into *M. leidyi* embryos by microinjection (Figure 10A). Similar to *Ml*Par-6-mVenus mRNA overexpression, the *Ml*Par-1-mCherry translated protein was observed after 4 hours post injection into the uncleaved egg (early localization in blastomeres was too faint to detect by this method). Our *in vivo* observations confirm the localization pattern described above by using *Ml*Par-1 antibody at gastrula stages. *Ml*Par-1-mCherry localizes uniformly and form aggregates in the cytosol during gastrulation (4-5 hpf; Figure 10A). This localization pattern remains throughout all recorded stages until cydippid juvenile stages where *Ml*Par-1-mCherry remains cytosolic in all cells but is highly concentrated in the tentacle apparatus and underneath the endodermal canals (24 hpf; Figure 10A and Movie 2).

**Figure 10.**
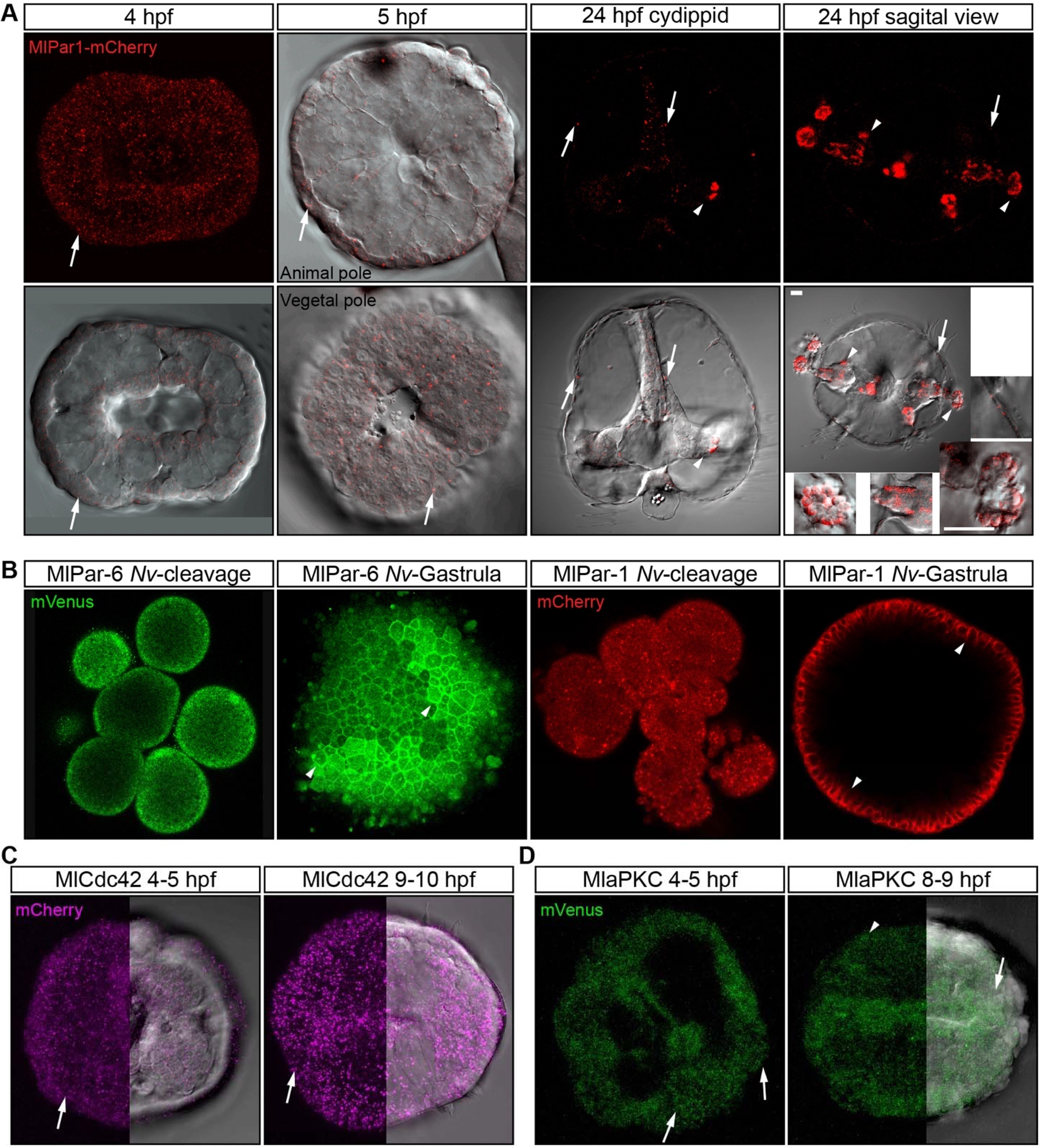
*In vivo* localization of *Ml*Par proteins during different stages of *M. leidyi* development. A) The overexpression of *Ml*Par1-mCherry protein displays similar patterns observed with the antibody staining against the same protein, with no cortical localization observed after 4 hpf. White arrows indicate *Ml*Par-1-mCherry protein cytosolic aggregates. White arrowhead indicates *Ml*Par-1-mCherry protein aggregates in the tentacle apparatus. All images are 3D reconstructions from a z-stack confocal series. Orientation of axes are depicted in the Figure. B) When ctenophore *Ml*Par6-mVenus and *Ml*Par1-mCherry are expressed in embryos of the cnidarian *N. vectensis*, the translated proteins display the same pattern than the previously described for endogenous *N. vectensis* proteins ^5^. White arrowheads indicate *Ml*Par6-mVenus and *Ml*Par1-mCherry cortical localization. All images are a single slices from a z-stack confocal series. C) *In vivo* localization of *Ml*Cdc42-mCherry. White arrows indicate *Ml*Cdc42-mCherry protein aggregates. All images are 3D reconstructions from a z-stack confocal series. D) *In vivo* localization of *Ml*aPKC-mVenus. White arrows indicate *Ml*aPKC-mVenus protein aggregates. All images are 3D reconstructions from a z-stack confocal series. Morphology is shown by DIC microscopy. See dynamic movements of these aggregates in Movie 1 and 2. Homogeneous cytosolic staining was digitally reduced to highlight cortical localizations.

*Ml*Par-1 aggregates may be consequence of the highly protein availability in the cytosol that is not captured to the cell cortex. Interestingly, the punctuate aggregates of *Ml*Par-1-mCherry are highly dynamic and move throughout the entire cytosol, suggesting a potential association with cytoskeletal components (see Movie 1). Unfortunately, we were not able to clone and express either *Ml*Dlg or *Ml*Lgl to observe if they resemble similar localization patterns to either *Ml*Par-1 or *Ml*Par-6 *in vivo*.

### MlPar-6 and MlPar-1 Structures are Conserved

To discount the possibility that the observations recorded *in vivo* for both *Ml*Par-6-mVenus and *Ml*Par-1-mCherry proteins are caused by a low-quality mRNA or lack of structural conservation, we overexpressed each ctenophore mRNA into embryos of the cnidarian *Nematostella vectensis* and followed their localization by *in vivo* imaging (Figure 10B). In these experiments, both *Ml*Par-6-mVenus and *Ml*Par-1-mCherry translated proteins display the same pattern than the previously described endogenous *N. vectensis* proteins do ^5^. In *N. vectensis* embryos, *Ml*Par-6-mVenus and *Ml*Par-1-mCherry symmetrically distribute during early cleavage stages and both protein asymmetric localization was observable only after blastula formation (Figure 10B). These data not only suggest that the ctenophore *Ml*Par-6 and *Ml*Par-1 protein structure and function are conserved but also indicate that their subcellular localization in *M. leidyi* embryos is regulated by an interaction with other signaling pathways that is different from the *N. vectensis* embryos.

### MlCdc42 and MlaPKC

To expand our observations on the localization of Par system components, we separately overexpressed the mRNA encoding for *Ml*Cdc42 fused to mCherry (*Ml*Cdc42-mCherry) and *Ml*aPKC fused to mVenus (*Ml*aPKC-mVenus). Both *Ml*Cdc42-mCherry (Figure 10C) and *Ml*aPKC-mVenus (Figure 10D) translated protein were observable and recorded after 3 to 4 hours post injection into the uncleaved egg. Surprisingly, *Ml*Cdc42-mCherry only localizes in the cytosol and form punctate aggregates without any clear cortical localization through all stages recorded (Figure 10C). On the other hand, *Ml*aPKC-mVenus displayed an apparent apical localization but the signal of the translated protein was not strong enough to discriminate this localization from the cytosolic protein (Figure 10D). Even though these data give some insights on the localization of both *Ml*Cdc42-mCherry and *Ml*aPKC-mVenus proteins, the development of specific antibodies is required to determine the localization of maternally loaded proteins and discard technical artifacts. Unfortunately, for technical reasons we were not able at this time to co-inject these proteins together in order to see their *in vivo* interactions during embryonic development.

## DISCUSSION

### Par protein asymmetry is established early but not maintained during M. leidyi embryogenesis

The asymmetric localization of the Par/aPKC complex has been used as an indicator of apical-basal cell polarity in a set of animals, including bilaterians^4,5,14,15,53,56–59,63,66,68,69,76–80^ and a cnidaria^5, 21^. While in the studied bilaterians this asymmetry is established and maintained since the earliest stages of development^60,63,65,71,88,89^, in the cnidarian *N. vectensis* there is no early asymmetrical localization of any of the Par components^5, 21^ and embryonic polarity is controlled by the Wnt signaling system^16,73,90–92^. In spite of these differences, once epithelial tissues form and true-epithelial cell-polarity is established in both bilaterian and cnidarian species, the asymmetric localization of Par proteins become highly polarized and is maintained through development. In those cases, Par-mediated apicobasal cell polarity is responsible for the maturation and maintenance of cell-cell adhesion in epithelial tissue^4, 22^. We have suggested that the polarizing activity of the Par system was already present in epithelial cells of the MRCA between Bilateria and Cnidaria^5,22^ and could be extended to all Metazoa where these proteins are present (including ctenophores, sponges, and placozoans^20,23^).

However, our current data suggest a different scenario for ctenophores where the Par protein polarization observed during earlier stages (characterized by the apical and cortical localization of *Ml*Par-6) is not maintained when ctenophore juvenile epithelial tissues form after 9 hpf. Embryos of the ctenophore *M. leidyi* are highly polarized^75–77,93–95^ and the asymmetrical localization of *Ml*Par-6 to the animal pole in the uncleaved zygote and the apical cortex of blastomeres (Figure 3) may be a consequence of this maternally contributed polarization. Even though this cortical polarization of *Ml*Par-6 is maintained during gastrulation in the ectoderm (Figures 4 and 5), epithelial cells of later cydippid stages do not display an asymmetric localization of *Ml*Par-6 (Figure 6). In addition, the subcellular localization of *Ml*Par-1 does not display a clear asymmetric localization during any of the observed developmental stages (Figures 7 to 9). Instead, punctate aggregates distribute symmetrically and actively migrate throughout the cytosol.

In bilaterians, the mechanism that induces and establishes the asymmetrical localization of Par proteins in the cell cortex during early animal development remains unknown, even though the maintenance of the apicobasal cell polarity in mature epithelial tissue is carried out by a conserved and well-studied mechanism. While the Par/aPKC complex is localized to the apical cortex by its interaction with the CCC and the Crb complex^4,20,23,96,97^, the cortico-lateral localization of Par-1 depends on the stable interaction between SJs and the Scribble complex^20,23,44,46–49,98^. Mutations in any of these components in bilaterian embryos results in a mislocalization of polarity proteins and the disassembling of junctional complexes^4,22,45,46,48,49,51–53^.

The components of the ctenophore *Ml*Par/aPKC complex (*Ml*Par-3/*Ml*aPKC/*Ml*Par-6 and *Ml*Cdc42) are highly conserved and contain all the domains present in other metazoans (Supplementary Figure 4)^20, 23^. Therefore, they are not only able to phosphorylate and displace *Ml*Par-1 and *Ml*Lgl to the cytoplasm but also are able to interact with Crb and localize to the apical cortex (Figure 10B). Similarly, the primary structure of *Ml*Par-1 protein (a Serine/threonine-protein kinase) is highly conserved and contains all the domains (with the same amino acid length) required for its proper functioning in other metazoans (Supplementary Figure 4)^20, 23^, and localizes to the lateral cortex when expressed in cnidarian embryos (Figure 10B). Regardless, these proteins do not asymmetrically localize to the cortex of *M. leidyi* juvenile epithelium.

Recent studies have shown that ctenophores do not have homologous for any of the Crb complex components^23^, required for the proper stabilization of the CCC and Par/aPKC complex in other studied taxa^4,33,37,64,99,100^. The lack of *Ml*Par-6 (Figure 5) and *Ml*Cdc42 (Figure 8C) polarization during later stages is totally congruent with these observations, indicating that Par proteins in ctenophores do not have the necessary interactions to stabilize apico-basal cell polarity in their cells as in other animals. In addition, ctenophore species do not have the molecular components to form SJs and lack a Scribble homolog^23, 48^. This suggests that none of the lateral polarity proteins (Dlg, Lgl, and Par-1) can localize to the lateral cortex of the ctenophore cells and explains the cytosolic localization of *Ml*Par-1 during the observed stages^42,45,49,101,102^.

Furthermore, it would be interesting to asses if the asymmetric localization of *Ml*Par-6 in early *M. leidyi* embryos, like in bilaterians, has roles related with the orientation of cell divisions and the establishment of cell-cell adhesion. The conservation of *Ml*Par-6 protein structure and apical localization during early cleavage stages also means that *Ml*Par-6 might be able to interact with cadherin, but that the Crb complex is not required for its polarization^103^. Unless the embryonic cell polarity in ctenophores is mediated by a completely different mechanism than bilaterians (a highly likely possibility because it is not clear that cnidarians use bilterian-spefic mechanism either), our data would imply that cnidarians, such as *N. vectensis*, gained a mechanism that maintains apicobasal cell polarity in epithelial tissues that was retained in the bilaterian radiation.

Unfortunately, due to technical difficulties we are unable yet to test the function of maternally loaded *Ml*Par proteins during the early development of *M. leidyi*. To solve this question, further research in a larger sample of non-bilaterians animals is also required.

### Evolution of cell polarity and epithelial structure in Metazoa

Given the genomic conservation of cell-polarity components in the Bilateria and Cnidaria, we propose to re-define a ‘true’ epithelium (classical bilaterian definition) to include its mechanistic regulatory properties. That is, the structural properties of a ‘true’ epithelium are the result of conserved interactions between subcellular pathways that polarize epithelial cells. Thus, when we seek to understand the origins of the epithelial nature of one particular tissue, we are trying to understand the synapomorphies (shared derived characters) of the mechanisms underlying the origin of that particular tissue. Under this definition, a ‘true-epithelium’ may have a single origin in Metazoa, but, different mechanisms might be co-opted to generate similar epithelial morphologies. Ctenophore epithelia, along with other recent work in *N. vectensis* endomesoderm^5, 22^ and *Drosophila* midgut^103^, suggest this possibility. In all these cases, epithelial cells are highly polarized along the apical-basal axis, but this polarization does not depend on Par proteins. Therefore, these cells are not able to organize a ‘true-epithelium’ (mechanistic definition) and the homology of this tissue cannot be extended to all Eumetazoa.

Recent genomic studies have shown that, during animal evolution, different signaling pathways components have emerged at different nodes of the metazoan tree^23,26,28^. Interestingly, ctenophore genomes suggest that new signaling pathways do not only emerge as individual components but also as a full set of protein complexes that bring a new set of interactions that remodel the cell structure and behavior. This reinforces the idea that signaling pathways act as modules that interact with each other to organize epithelial cells^104–106^.

Genomic studies also suggest that in addition to the origin of complex signaling pathways, ctenophore species lack the molecular interactions necessaries to form the apical cell polarity and junctions observed in Cnidaria + Bilateria. Intriguingly, ctenophore genomes do not have the Wnt signaling pathway components^26,27,107^ that control the activity of Par proteins in bilaterian and cnidarian embryos (components that are also present in poriferan and placozoan genomes^23^). For example, in bilaterians the Wnt/PCP signaling pathway antagonizes the action of the Par/aPKC complex^7,10,15,49,108,109^, so this may explain the lack of polarization in ctenophore tissue. Furthermore, ctenophore species do not have mature AJs and the full set of cell-cell adhesion proteins^23,26,48^ as we know them in other metazoans, including Placozoans and Poriferans^1, 23^. The cadherin of ctenophores does not have the cytoplasmic domains required to bind any of the catenins of the CCC (e.g. p120, alpha- and ß-catenin)^23^. This implies that neither the actin nor microtubule cytoskeleton can be linked to ctenophore cadherin through the CCC, as seen essential in other metazoans to polarize Par proteins. This suggests that there are additional mechanisms that integrate the cytoskeleton of ctenophore cells with their cell-cell adhesion system. A recent work^110^ claims that other species of ctenophore (*Beroe* and *Pleurobrachia*), but not *Mnemiopsis leidyi*, have basement membranes underlying adult tissue; however, all three species whose genomes have been sequenced appear to have the same complement of ECM components (e.g. Fig. 2^110^).

Because *Mnemiopsis* clearly have polarized epithelia^111^ (Supplementary Figure 5), it is likely the differences in lobate ctenophores is a fixation artifact at the stage or tissue the material was prepared (these data were not mentioned^110^). In conclusion, we have shown that regardless of the high structural conservation of Par proteins across Metazoa, ctenophore cells do not have other essential components to deploy and/or stabilize the asymmetrical polarizing function of the Par system as in other studied metazoans. Thus, ctenophore tissues organize their epithelium in a different way than the classical definition seen in Bilaterians. In agreement with genomic studies, our results question what molecular properties defined the ancestral roots of metazoan epithelium, and whether similar epithelial morphologies (e.g., epidermis and mesoderm) could be developed by independent or modifications of existing cellular and molecular interactions (including cell adhesion systems). Unless the lack of Par protein localization in *M. leidyi* is a secondary loss, the absence of these pathways in ctenophores implies that a new set of interactions emerged at least in the Cnidaria+Bilateria ancestor (Figure 11), and that, could have regulated the way by which the Par system polarizes embryonic and epithelial cells. While bioinformatic studies are critical to understand the molecular composition, we need further research to understand how these molecules actually interact with one another to organize cellular behavior (e.g., integrin-collagen, basal-apical interactions) in a broader phylogenetical sample, including Porifera and Placozoa.

**Figure 11.**
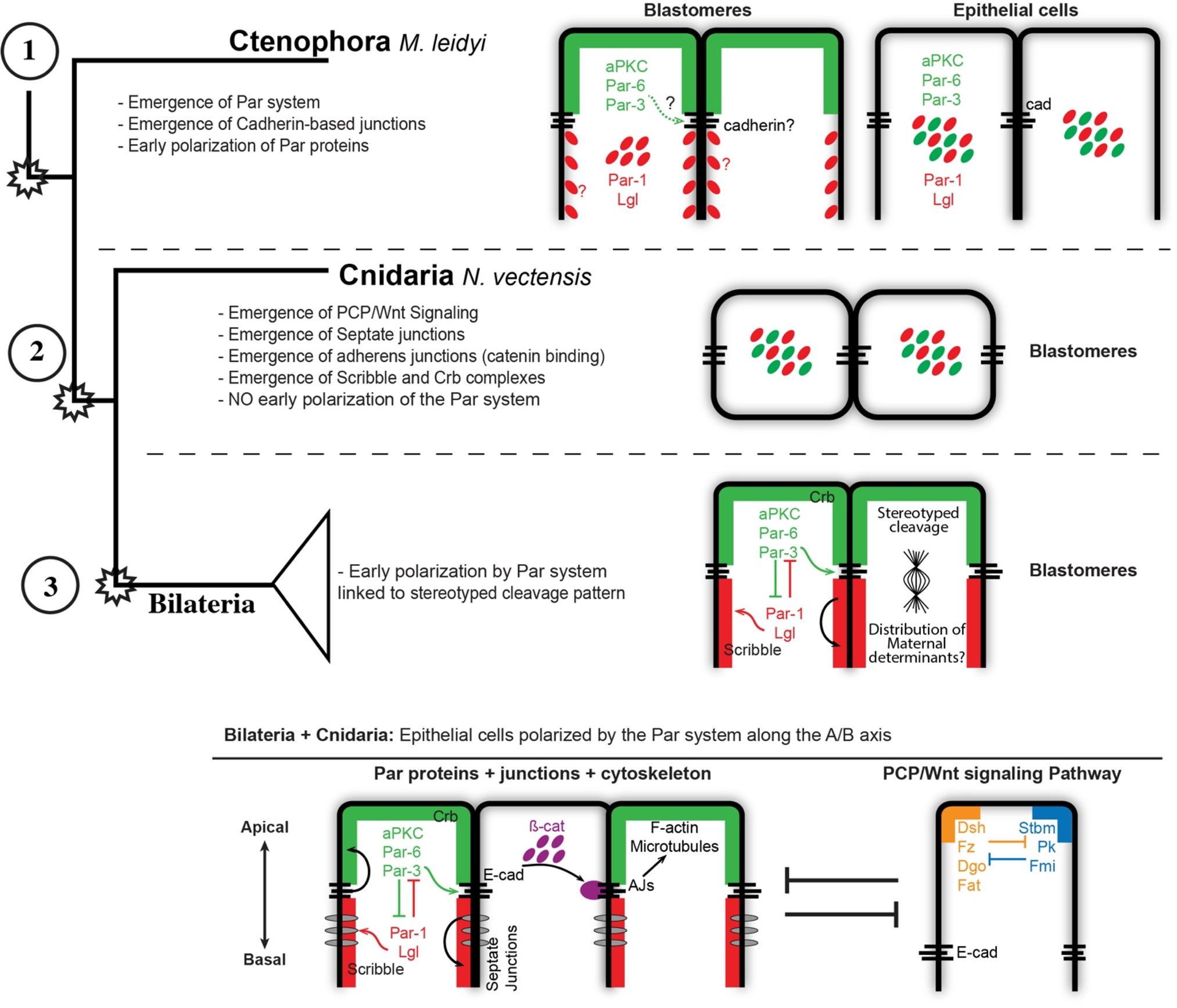
Evolution of cell polarity in Metazoa. Diagram depicting the evolution of different interactions between known signaling pathways that organize cell polarity in animal cells, including the new information obtained by this study during the early development of the ctenophore *M. leidyi*. Our results challenge the conception of a deep homology of the true-epithelial structure and the establishment of the apicobasal cell polarity in Metazoa.

## METHODS

### Culture and Spawning of M. leidyi

Spawning, gamete preparation, fertilization and embryo culturing of *M. leidyi* embryos was performed as previously described^112^. Adult *M. leidyi* were maintained at the Whitney Laboratory for Marine Bioscience of the University of Florida (USA) in constant light conditions. Spawning was induced by incubating the adults in a three to four-hour dark cycle at room temperature (approximately 25°C). The embryos were washed multiple times in filtered SW and if dechorionated, were kept in glass or gelatin coated dishes (to prevent sticking) at room temperature until the desired stage.

### Western Blot

Western blots were carried out as described^5, 22^ using adult epithelial tissue lysates dissected by hand in order to discard larger amount of mesoglea. Antibody concentrations for Western blot were 1:1,000 for all antibodies tested.

### Immunohistochemistry

All immunohistochemistry experiments were carried out using the previous protocol for *M. leidyi*^112^. Embryos were fixed on a rocking platform at room temperature. Embryos of different stages were fixed for 2 hours in fresh fixative (100mM HEPES pH 6.9; 0.05M EGTA; 5mM MgSO4; 200mM NaCl; 1x PBS; 3.7% Formaldehyde; 0.2% Glutaraldehyde; 0.2% Triton X-100; and 1X fresh sea water). Fixed embryos were rinsed at least five times in PBT (PBS buffer plus 0.1% BSA and 0.2% Triton X-100) for a total period of 3 hours. PBT was replaced with 5% normal goat serum (NGS) diluted in PBT and fixed embryos were blocked for 1 to 2 hours at room temperature with gentle rocking. Primary antibodies were diluted in 5% NGS-PBT to the desired concentration. Blocking solution was removed and replaced with primary antibodies diluted in NGS. All antibodies incubations were conducted over night on a rocker at 4°C. After incubation of the primary antibodies, samples were washed at least five times with PBT for a total period of 3 hours. Secondary antibodies were then applied (1:250 in 5% NGS-PBT) and samples were left on a rocker overnight at 4°C. Samples were then washed with PBT and left on a rocker at room temperature for an hour. Samples were then washed once with PBT and incubated with DAPI (0.1µg/µl in PBT; Invitrogen, Inc. Cat. # D1306) for 1 hour to allow nuclear visualization. Stained samples were rinsed again in PBS two times and dehydrated quickly into isopropanol using the gradient 50%, 75%, 90%, and 100%, and then mounted in Murray’s mounting media (MMM; 1:2 benzyl benzoate:benzyl alcohol) for visualization. Note that MMM attenuates the DAPI signal from samples. We scored more than 1,000 embryos per each antibody staining and confocal imaged more than 50 embryos at each stage obtaining similar staining patterns for each case. After 10-11 hpf, we scored larvae at 12, 15, and 24 hpf.

The primary antibodies and concentrations used were: mouse anti-alpha tubulin (1:500; Sigma-Aldrich, Inc. Cat.# T9026. RRID:AB_477593). Secondary antibodies are listed in the Key Resources table.

Rabbit anti-*Ml*Par-6, and rabbit anti-*Ml*Par-1 antibodies were custom made high affinity-purified peptide antibodies that commercially generated by Bethyl labs, Inc. (Montgomery, TX, USA). Affinity-purified *M. leidyi* anti-Par-6 (anti-*Ml*Par-6) and anti-Par-1 (anti-*Ml*Par-1) peptide antibodies were raised against a selected amino acid region of the *Ml*Par-6 protein (MTYPDDSNGGSGR) and *Ml*Par-1 protein (KDIAVNIANELRL), respectively. Blast searches against the *M. leidyi* genome sequences showed that the amino acid sequences were not present in any predicted *M. leidyi* proteins other than the expected protein. Both antibodies are specific to *M. leidyi* proteins (Figure 2) and were diluted 1:100.

### mRNA Microinjections

The coding region for each gene of interest was PCR-amplified using cDNA from *M. leidyi* embryos and cloned into pSPE3-mVenus or pSPE3-mCherry using the Gateway system^113^. To confirm the presence of the transcripts during *M. leidyi* development, we cloned each gene at 2 hpf and 48 hpf. Eggs were injected directly after fertilization as previously described for *N. vectensis* studies^5,114,115^ with the mRNA encoding one or more proteins fused in frame with reporter fluorescent protein (N-terminal tag) using an optimized final concentration of 300 ng/µl for each gene. Fluorescent dextran was also co-injected to visualize the embryos. Live embryos were kept at room temperature and visualized after the mRNA of the FP was translated into protein (4-5 hours). Live embryos were mounted in 1x sea water for visualization. Images were documented at different stages. We injected and recorded at least 20 embryos for each injected protein and confocal imaged each specimen at different stages for detailed analysis of phenotypes in vivo. We repeated each experiment at least five times obtaining similar results for each case. The fluorescent dextran and primers for the cloned genes are listed in Key resources table.

### Imaging of *M. leidyi* Embryos

Images of live and fixed embryos were taken using a confocal Zeiss LSM 710 microscope using a Zeiss C-Apochromat 40x water immersion objective (N.A. 1.20). Pinhole settings varied between 1.2-1.4 A.U. according to the experiment. The same settings were used for each individual experiment to compare control and experimental conditions. Z-stack images were processed using Imaris 7.6.4 (Bitplane Inc.) software for three-dimensional reconstructions and FIJI for single slice and movies. Final figures were assembled using Adobe Illustrator and Adobe Photoshop.

Par proteins display a general cytosolic localization when their polarizing activity is inactive. This signal was diminished by modifying contrast and brightness of the images in order to enlighten their cortical localization (active state in cell-polarity and stronger antibody signal) as it has shown in other organisms. All RAW images are available upon request.

## ACKNOWLEDGMENTS

We thank C.E. Schnitzler and J. Ryan for technical assistance with the ctenophore genome. This research was supported by the NSF IOS-1755364 and NASA 16-EXO16_2-0041 from MQM.

## AUTHOR CONTRIBUTIONS

MS-S., and MQM. designed research and analyzed data. MS-S. performed research with help of MQM. MS-S., and MQM. wrote the manuscript. All authors read and approved the final manuscript.

---

Movie 1. Punctuate aggregates of *Ml*Par-1-mCherry are highly dynamic. 2.5 minutes *in vivo* recording of a a gastrula embryo at 40x.

Movie 2. Z-stack of *Ml*Par-1-mCherry expression at 24 hpf at 40X.

**Supplementary Figure 1.**
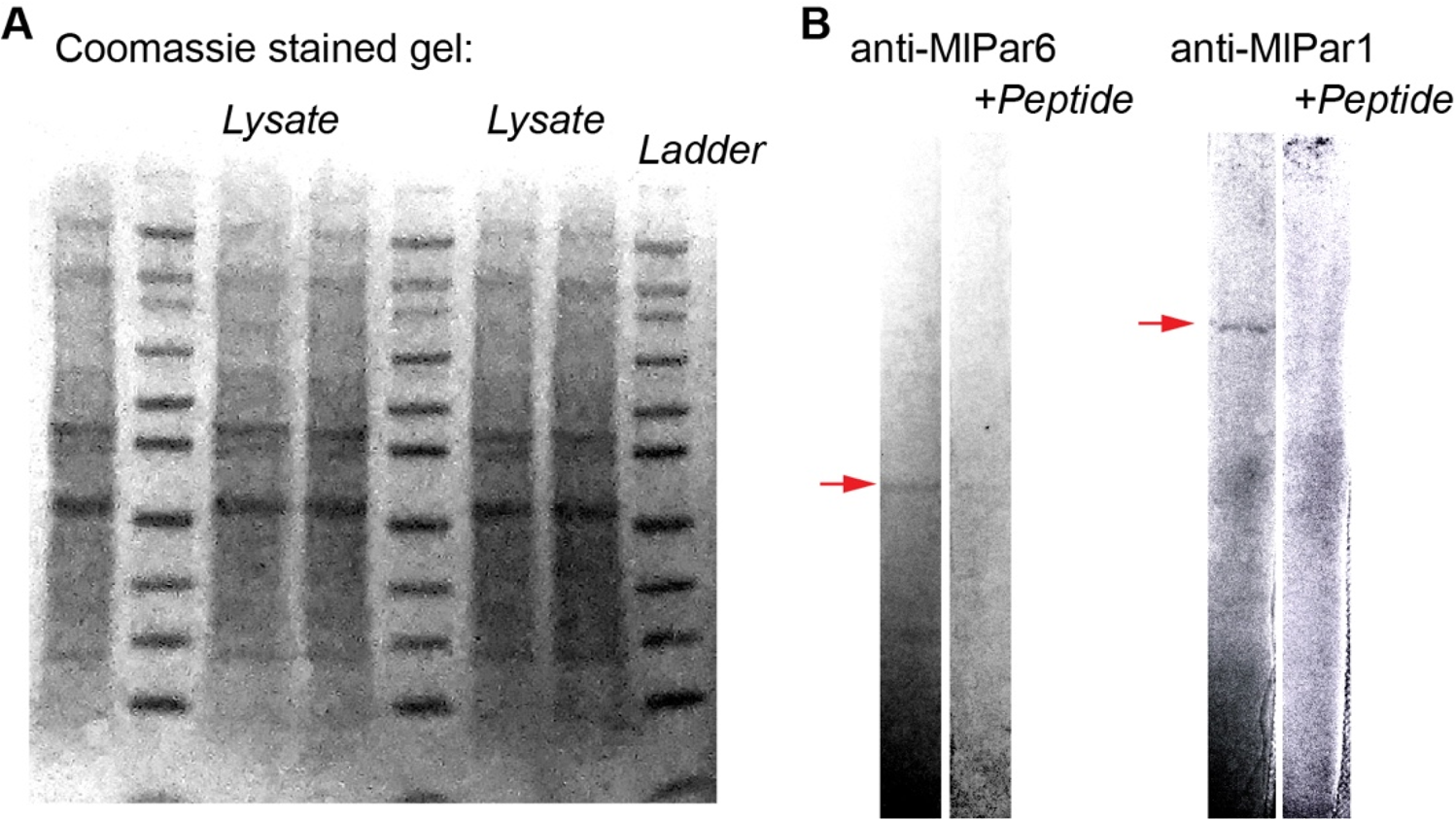
Supporting Figure related to Figure 2. A) Coomassie stained gel of the full input lysate. B) Full Western blot lanes for the MlPar-6 and MlPar-1 antibodies sections shown in Figure 2.

**Supplementary Figure 2.**
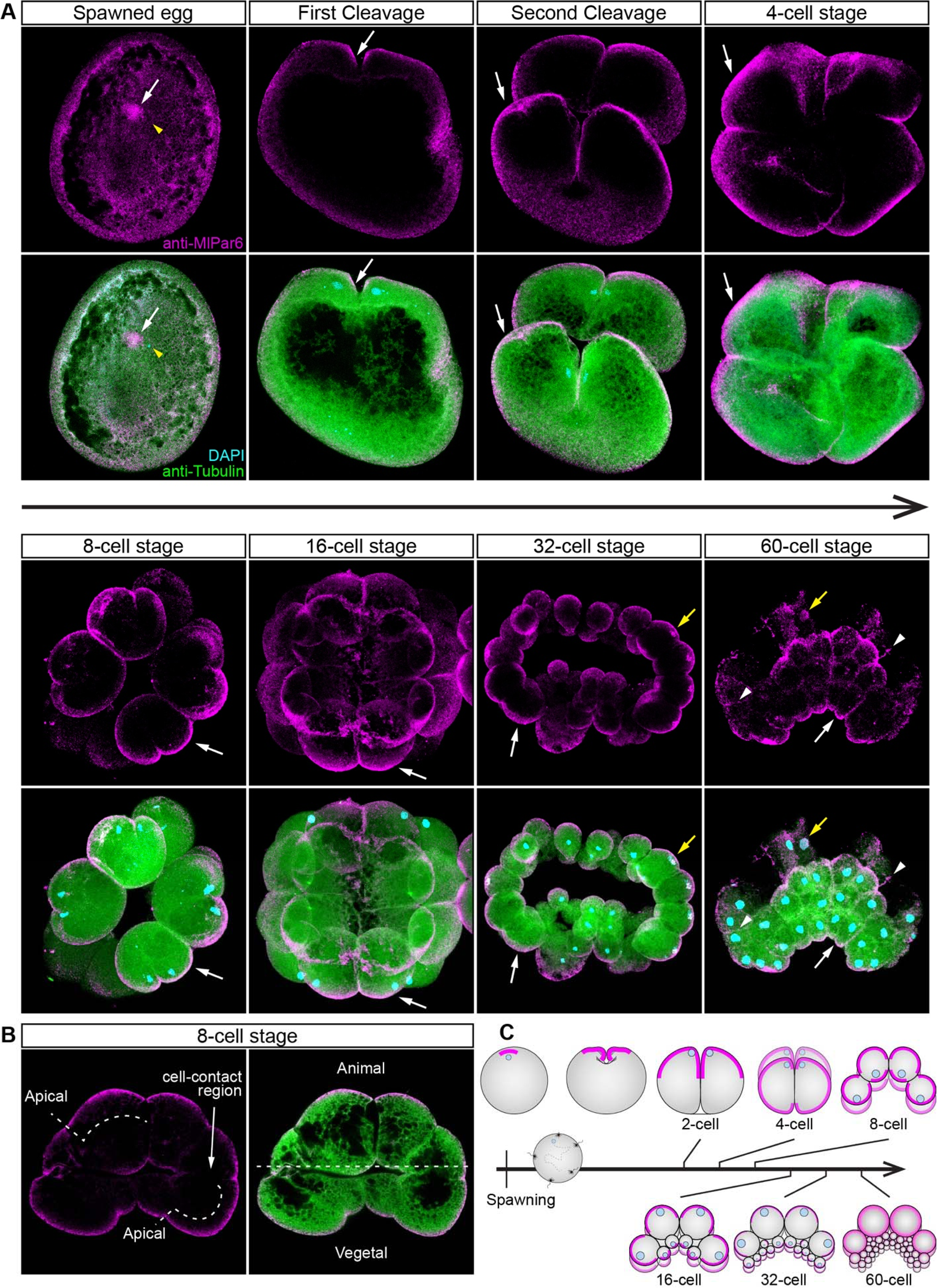
*Ml*Par-6 protein localizes asymmetrically in the cell cortex of the eggs and in the cell-contact-free regions of cleavage stages. A) Immunostaining against *Ml*Par-6 during cleavage stages of *M. leidyi* development. *Ml*Par-6 protein localizes to the apical cortex (white arrows) until the 60-cell stage where its signal was detected in regions of cell-contact (white arrowhead). Images are 3D reconstructions from a z-stack confocal series. B) A single optical section from the 8-cell stage z-stack confocal series shown in A. *Ml*Par-6 protein localizes to the apical cortex of the cells but is absent from cell-contact regions. Orientation of samples are indicated. C) Diagram depicting the cortical localization of *Ml*Par-6. The animal pole is to the top as depicted in B. Morphology is shown by DAPI and tubulin immunostainings. Yellow arrowhead indicates the zygote nucleus. Yellow arrows indicate nuclear localization. Homogeneous cytosolic staining was digitally reduced to highlight cortical localizations.

**Supplementary Figure 3.**
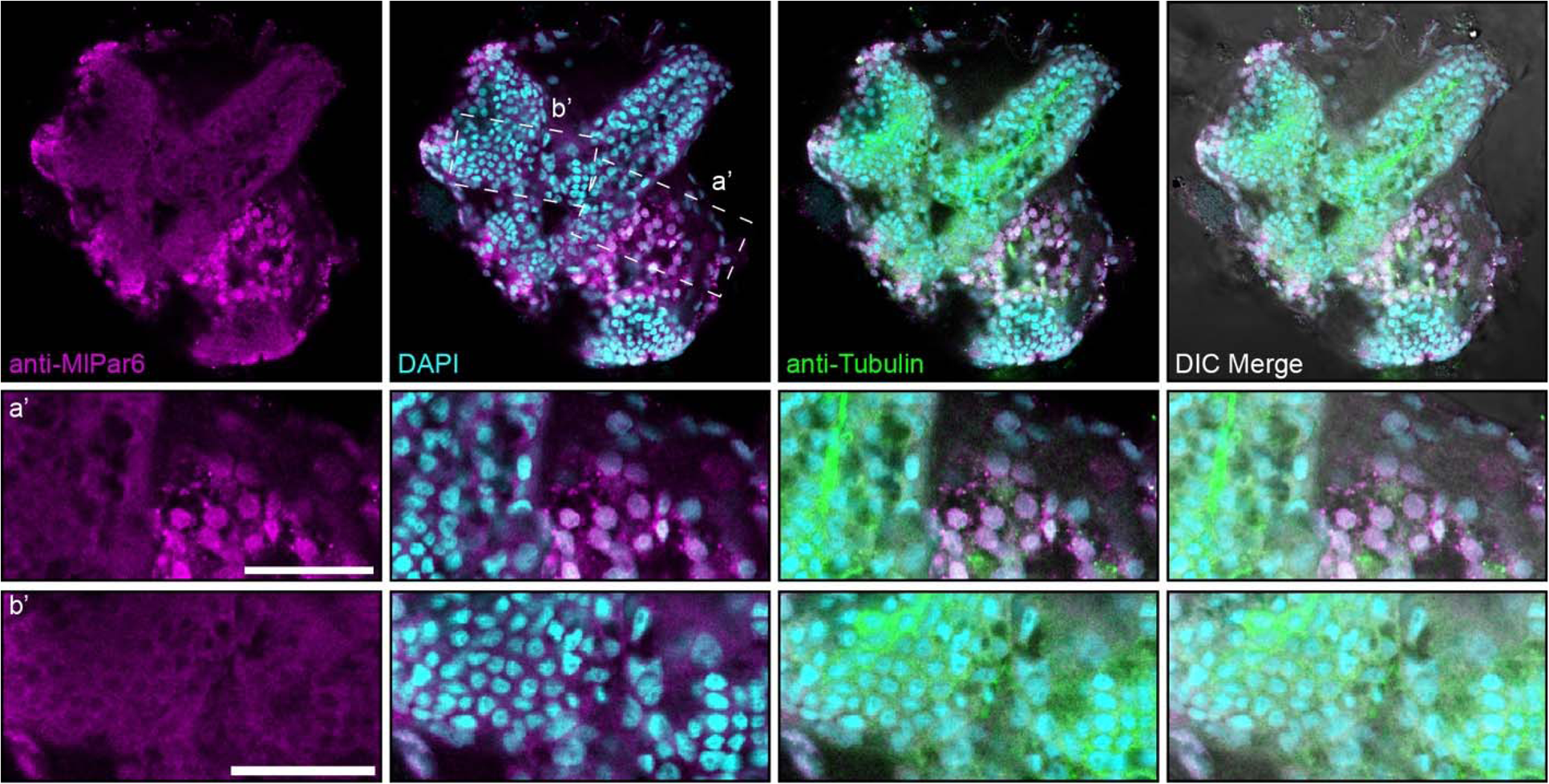
Immunofluorescent staining against MlPar-6 after 20 hpf. The distribution of *Ml*Par-6 in juvenile epithelium is nuclear and cytosolic during later stages and no asymmetrical and cortical localization was observed. a’ and b’ correspond to the magnifications of the regions depicted. Morphology is shown by DIC, DAPI, and tubulin immunostainings. Scale bars: 20 µm.

**Supplementary Figure 4.**
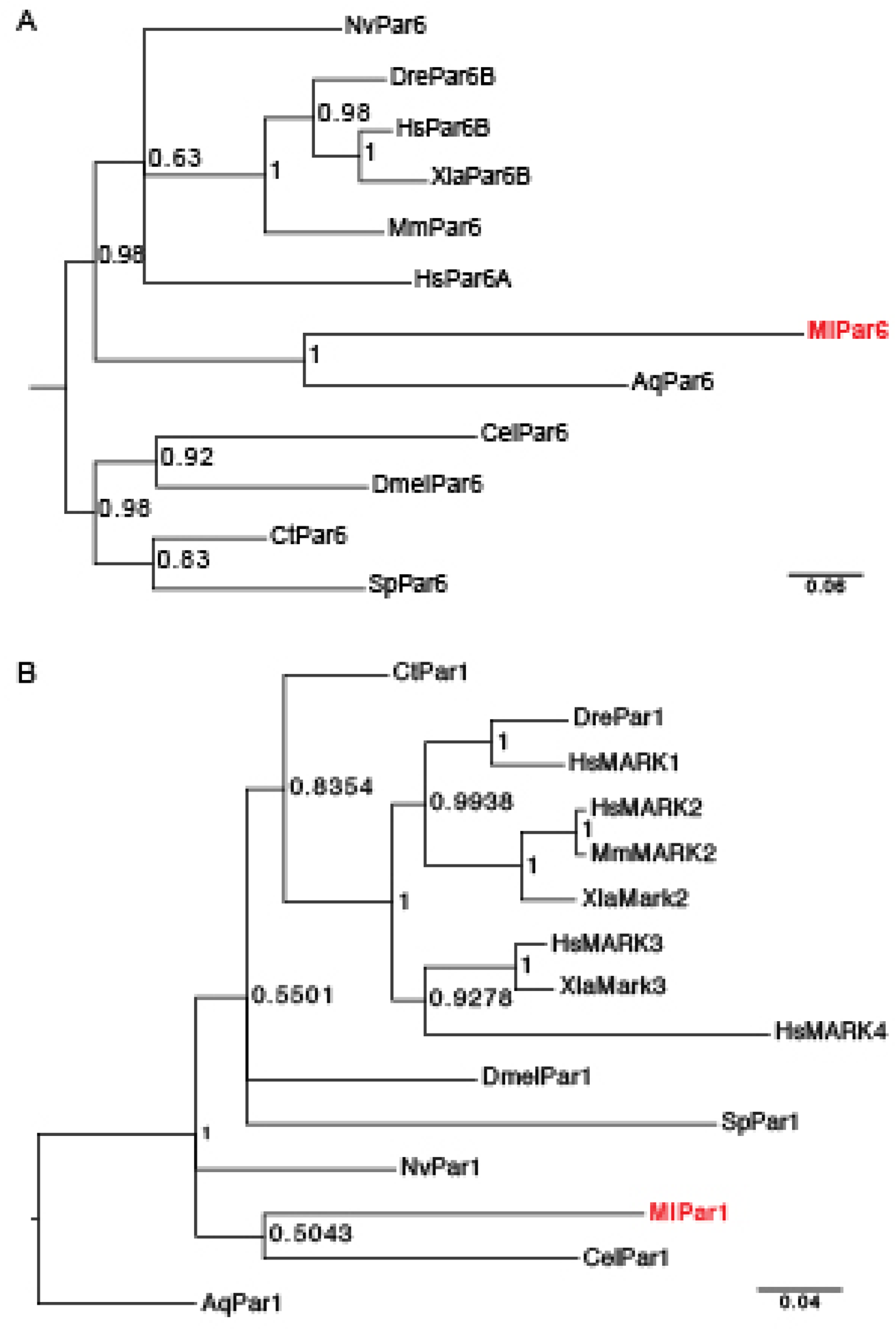
Phylogenetic analysis for (A) *Ml*Par-6 and (B) *Ml*Par-1. Trees were constructed using MrBayes (v3.2.6 x64) and consisted of 2,000,000 generations using “mixed” models. Maximum-likelihood tree bootstraps were based on 100 replicates. Aq: *Amphimedon queenslandica*; Cel: *Caenorhabditis elegans*; Ct: *Capitella teleta*; Dmel: *Drosophila melanogaster*; Dre: *Danio rerio*; Hs: *Homo sapiens*; Ml: *Mnemiopsis leidyi*; Mm: *Mus musculus*; Nv: *Nematostella Vec*tensis; Sp: *Strongylocentrotus purpuratus*; Xla: *Xenopus laevis*. See also references 20 and 23.

**Supplementary Figure 5.**
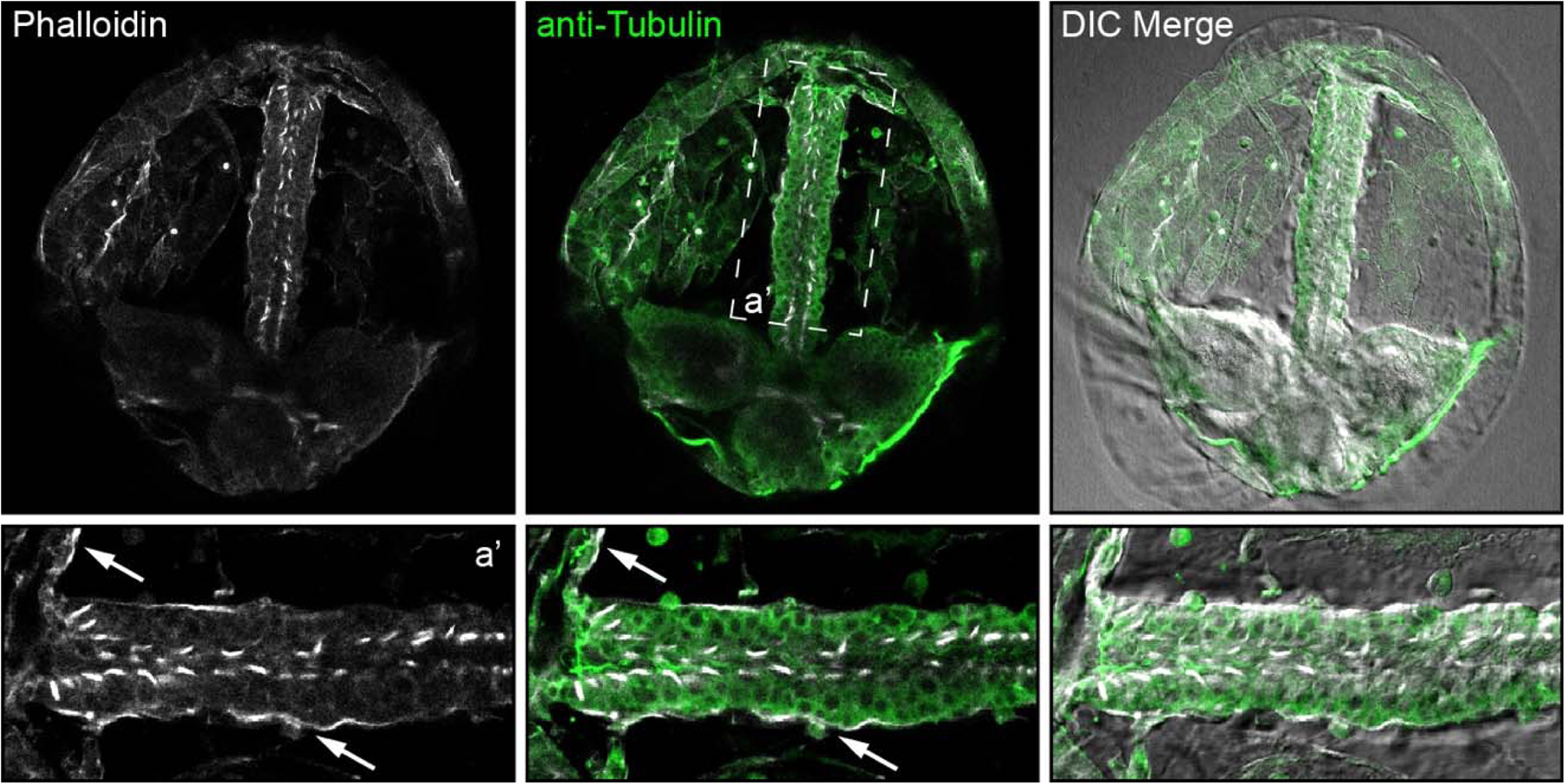
Phalloidin staining of *M. leidyi* cydippid suggest the presence of a basement membrane in this species. White arrows show the basement of the epithelium in contact with the ECM and mesoglea. a’ corresponds to the magnification of the depicted area (oral is at the left). Morphology is shown by tubulin immunostaining and DIC.

## Key resources table

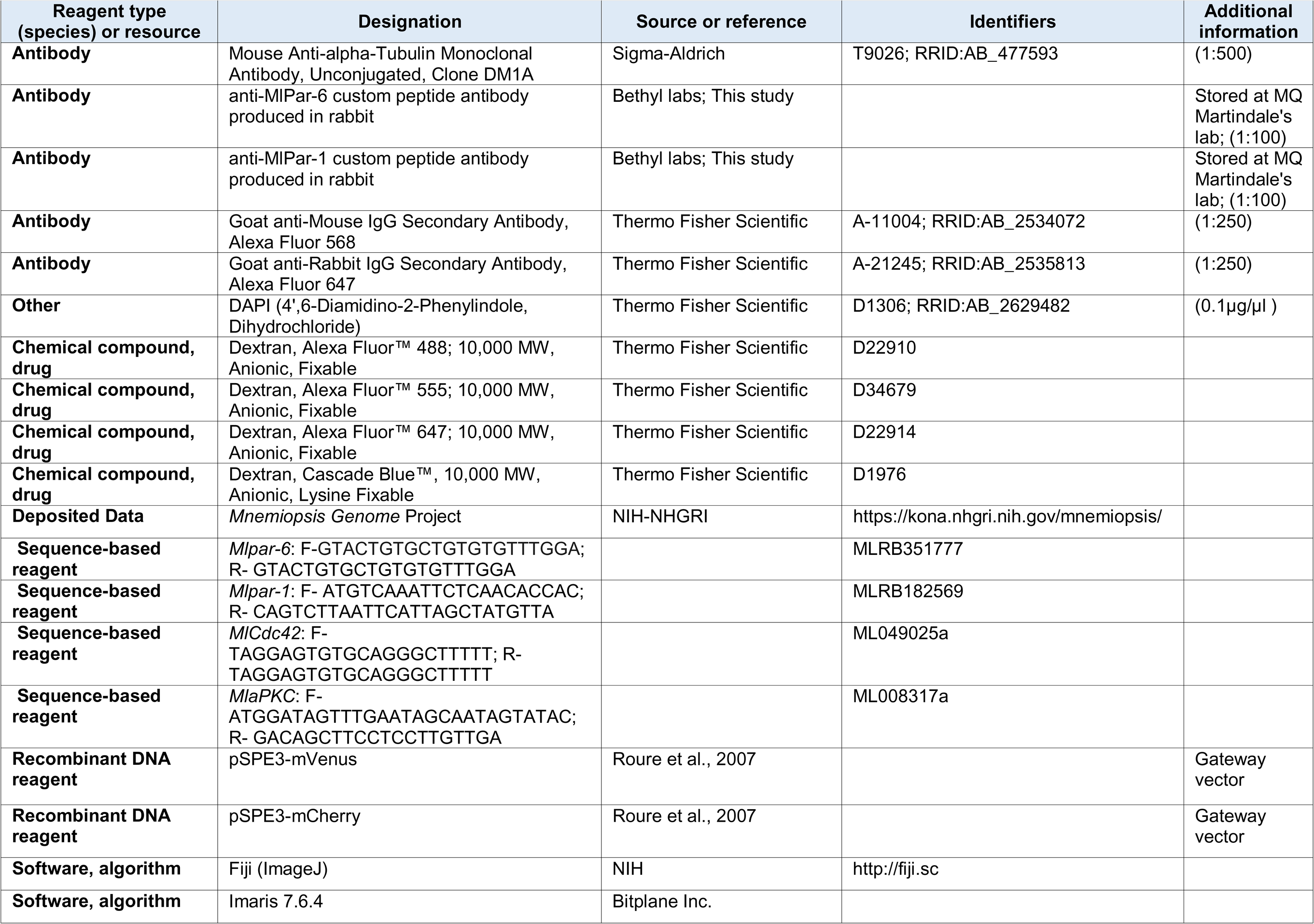

